# Can Single Cell Respiration be Measured by Scanning Electrochemical Microscopy (SECM)?

**DOI:** 10.1101/2023.04.24.538172

**Authors:** Kelsey Cremin, Gabriel N. Meloni, Dimitrios Valavanis, Orkun S. Soyer, Patrick R. Unwin

## Abstract

Ultramicroelectrode (UME), or - equivalently - microelectrode, probes are increasingly used for single-cell measurements of cellular properties and processes, including physiological activity, such as metabolic fluxes and respiration rates. Major challenges for the sensitivity of such measurements include: (i) the relative magnitude of cellular and UME fluxes (manifested in the current); and (ii) issues around the stability of the UME response over time. To explore the extent to which these factors impact the precision of electrochemical cellular measurements, we undertake a systematic analysis of measurement conditions and experimental parameters for determining single cell respiration rates, via the oxygen consumption rate (OCR) at single HeLa cells. Using scanning electrochemical microscopy (SECM), with a platinum UME as the probe, we employ a self-referencing measurement protocol, rarely employed in SECM, whereby the UME is repeatedly approached from bulk solution to a cell, and a short pulse to oxygen reduction reaction (ORR) potentials is performed near the cell and in bulk solution. This approach enables the periodic tracking of the bulk UME response to which the near-cell response is repeatedly compared (referenced), and also ensures that the ORR near the cell is performed only briefly, minimizing the effect of the electrochemical process on the cell. SECM experiments are combined with a finite element method (FEM) modeling framework, to simulate oxygen diffusion and the UME response. Taking a realistic range of single cell OCR to be 1×10^−18^ to 1×10^−16^ mol s^-1^, results from the combination of FEM simulations and self-referencing SECM measurements show that these OCR values are at - or below - the present detection sensitivity of the technique. We provide a set of model-based suggestions for improving these measurements in the future, but highlight that extraordinary improvements in the stability and precision of SECM measurements will be required if single cell OCR measurements are to be realized.

**TOC:** 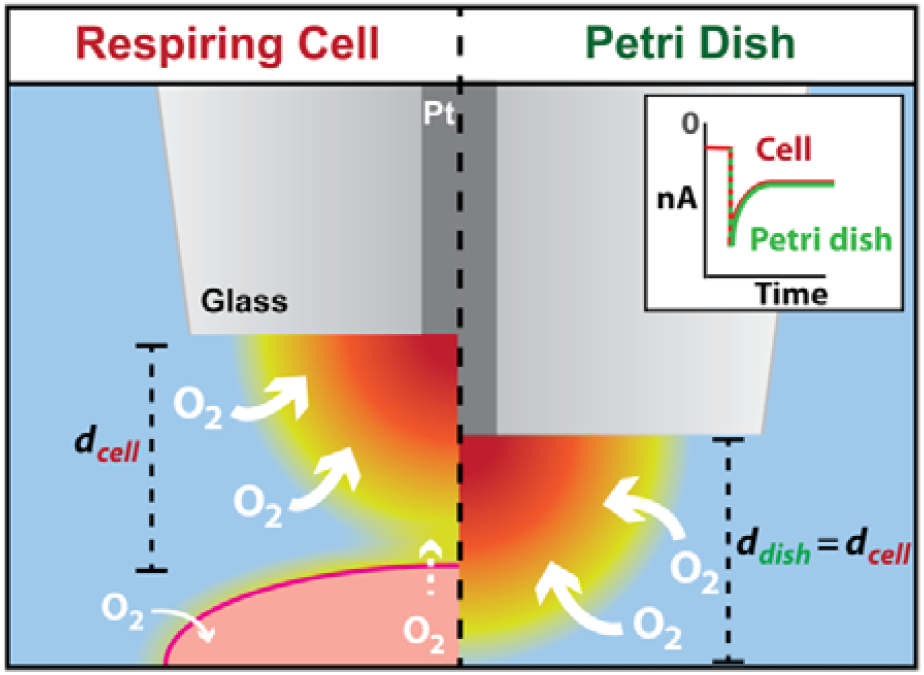

## Introduction

Precise measurement of cellular respiration rates is crucial for understanding the metabolic behavior of cells.^1^ Changes to respiration rates when cells are challenged with different conditions can aid in the elucidation of the overall cell metabolism. For example, depending on the experimental conditions, a reduced rate of respiration within cancerous cell lines could imply a shift to glycolytic and fermentative pathways, as described by the Warburg effect.^2, 3^

Respiration is usually quantified by bulk measurements of the oxygen consumption rate (OCR) of a population of cells. While these measurements can be accurate and sensitive,^4, 5^ they do not provide information on individual cell behaviors. Bulk measurements cannot be used to study interesting aspects, such as population heterogeneity in OCR or asynchronous glycolytic oscillations at the single cell level.^6^ Fluorescence microscopy offers a means to study individual cells within a population, and can be used to monitor respiration and oxidative stress, but is not reliably quantitative.^7^ Thus, quantitative single cell OCR measurements remain a significant experimental and instrumental challenge.

Ultramicroelectrodes (UMEs), or - equivalently - microelectrodes, are attractive for single cell measurements,^8–13^ particularly when used as the probe in scanning electrochemical microscopy (SECM),^14^ for which there are a diverse range of cellular studies.^15–18^ The coupling of fluorescence microscopy techniques, such as confocal laser scanning microscopy (CLSM), with SECM extends the depth of analysis of individual cells and related bilayer processes.^19–21^

Oxygen detection at UMEs is readily accomplished through the oxygen reduction reaction (ORR), which is a 4-electron process at platinum electrodes,^22–24^ with the resulting diffusion-limited cathodic current proportional to the local oxygen concentration.^25, 26^ When a UME is used as an SECM probe and is brought into the vicinity of a single cell, the response depends on the local oxygen concentration distribution (field), which is affected by the cell’s OCR. Several SECM studies have related ORR current to the OCR of cells and tissues.^1, 27–30^ Single cell OCR rates, approximated from population measurements, across a range of different commonly studied cell lines, range from *ca*. 5×10^−18^ to 5 ×10^−17^ mol s^-1^.^31^ However, single-cell SECM measurements have yielded OCR values that are considerably higher than these population-level estimates, by up to two orders of magnitude.^25, 30, 32–34^ An important questions is: how does this appreciable difference between SECM-based measurements and population-based estimates arise? Furthermore, what is the limit of detection in SECM-based OCR measurements, and is it possible to use SECM to detect single-cell OCR at the value estimated from population-level measurements?

Here we employ a self-referencing SECM method to define the limitations of single-cell OCR measurements. Self-referencing SECM temporally modulates the position of a UME probe such that oxygen current measurements taken in the vicinity (and under the influence) of the cell are compared to those taken in the bulk solution.^35, 36^ Self-referencing has been successfully used for single cell flux measurements previously and is highlighted as a favorable approach to improve accuracy in SECM.^36–39^ By performing an identical measurement protocol with an UME positioned in the bulk position, then in the vicinity of a cell, and comparing the results, a self-referenced result series is created with increased sensitivity, particularly as it accounts (at least in part) for any change in the response in the UME during the course of a series of measurements. Alteration of the UME response is common in biological media,^33, 40, 41^ due to the proteins and other biological molecules which may coat and deactivate the electrode surface.^42–44^ Electrode surface fouling can lead to deterioration in the UME response and add a source of inaccuracy to measurements. The self-referencing approach can reveal the extent of deactivation in the measurements, by tracking the bulk response during the course of the measurements.

A further benefit of self-referencing SECM is that the electrode is only positioned near the cell for short periods of time, minimizing its impact on the cell, for example, from the hindrance of substrate (O_2_) transport into the gap between the UME and cell. Furthermore, by pulsing the potential to perform oxygen detection, for a relatively brief period, any effects arising from UME electrolysis are also minimized.

Here, we combine self-referencing SECM with extensive finite element method (FEM) modeling to explore single cell OCR measurements with the mammalian HeLa cell line. We find that the ability to measure single cell respiration via SECM depends crucially on several experimental factors, including UME geometry, how close the UME can be positioned near to a cell, as well as cell behavior. Our integrated modeling and experimental data show that when considering all these factors, single cell OCR measurements are very difficult to realize and could be easily misinterpreted. Thus, these findings highlight current challenges and limitations of SECM-based single cell OCR measurements and provide suggestions for future method development and improvements in electrochemical probe-based measurement systems.

## Methods

We provide details of experimental methods pertaining to SECM measurements and a brief explanation of FEM modeling. The *Supporting Information* (SI) contains further details of the cell culture (Section SI-1), UME fabrication and SECM platform instrumentation (Section SI-2), optical microscopy (Section SI-3) and FEM simulations (Section SI-4).

### SECM protocol

SECM was performed atop an inverted CLSM (Leica TCS SP5 X microscope). A two-electrode setup was employed with a 5 μm radius (*a*) platinum disk UME as the working electrode with a ratio of platinum electrode to overall probe radius (RG ratio) of 15, and a chloridized silver wire quasi-reference counter electrode (QRCE).^42^ The UME radius, before platinization (section SI-2), was calculated from cyclic voltammograms^45^ recorded with hexaammineruthenium(III) chloride in potassium chloride solution and from optical microscopy. The RG ratio was also determined from optical images. The SECM probe was mounted on a positioning stage controlled by a piezo manipulator (PI, P-611.3S XYZ Nanocube, 100 μm). With the aid of CLSM visualization, the SECM probe was positioned 10 μm above the top plane of a cell (UME-cell separation – *d*), then retracted, using the piezo actuator, to 100 μm above (approximately the bulk solution). A diagram of the overall measurement setup is shown in Figure 1A.

**Figure 1.**
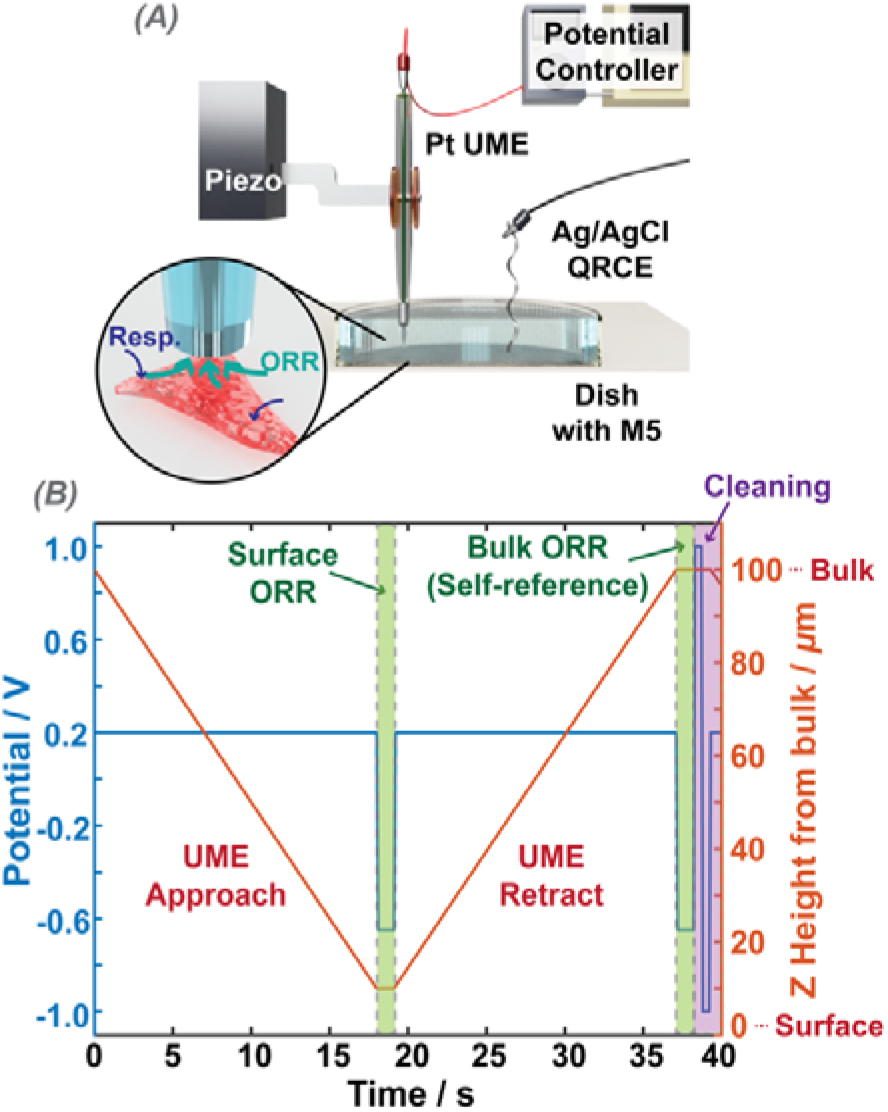
(A) Illustration of the SECM experimental set up. (B) Temporal profile of a single complete cycle of the (repeated) self-referencing program (one ‘hop’), where the potential program applied at the UME is shown on the left axis (blue), and the height position of the UME, measured from the top plane of the cell, is shown on the right axis (red).

Figure 1B summarizes the self-referencing protocol employed for the single cell OCR measurements, detailing the UME positional translation and potential control. A custom-built LabVIEW program (2019 release, National Instruments) was used to translate the UME, control the potential, and read the currents at the UME. First, the UME was brought towards the cell surface (speed of 5 μm s^-1^), while kept at a potential of 0.2 V vs. the QRCE, where there was no ORR. Upon reaching the pre-determined position about 10 μm above the top plane of the cell, the UME bias was switched to the ORR potential (-0.9 V) and the current recorded. The probe potential was then switched back to 0.2 V and the UME was moved back to the bulk position, where the ORR potential was applied again, and the current recorded for an equivalent length of time. Unless otherwise stipulated, data were recorded for the 1 second pulse lengths and we report herein on the current at 1 second, where the UME response was close to steady-state. The current measured with the UME at the cell vicinity was normalized to the corresponding signal with the UME at the bulk position. After the bulk self-referencing pulse, and before moving the probe near the cell, a series of short pulses (4 pulses, 0.5 seconds each) between -1 and 1 V were applied to clean the UME surface. The entire process, as described, was repeated *n* times (typically 20-30 times) with a set interval between each measurement (typically 30 seconds), so to record data near the cell over a period of time.

### FEM Simulations

A two-dimensional, axisymmetric cylindrical simulation representing the UME and cell was constructed in COMSOL Multiphysics (v. 5.5) (see Figure 2). The model comprised of two domains: D1 representing the bulk media, and D2 representing a HeLa cell. The cell was modeled as a half ellipsoid approximate in size to a HeLa cell (radius: 9 μm, height: 2.5 μm) and the UME geometry was parameterized based on electrochemical and microscopy characterization (planar disk, *a* = 5 μm, RG = 15). A realistic intracellular environment was simulated by setting the interior cell (cytosol) oxygen concentration to 14.5 μM,^46^ with a cytoplasmic diffusion coefficient of 7×10^−11^ m^2^ s^-1^.^47^

**Figure 2.**
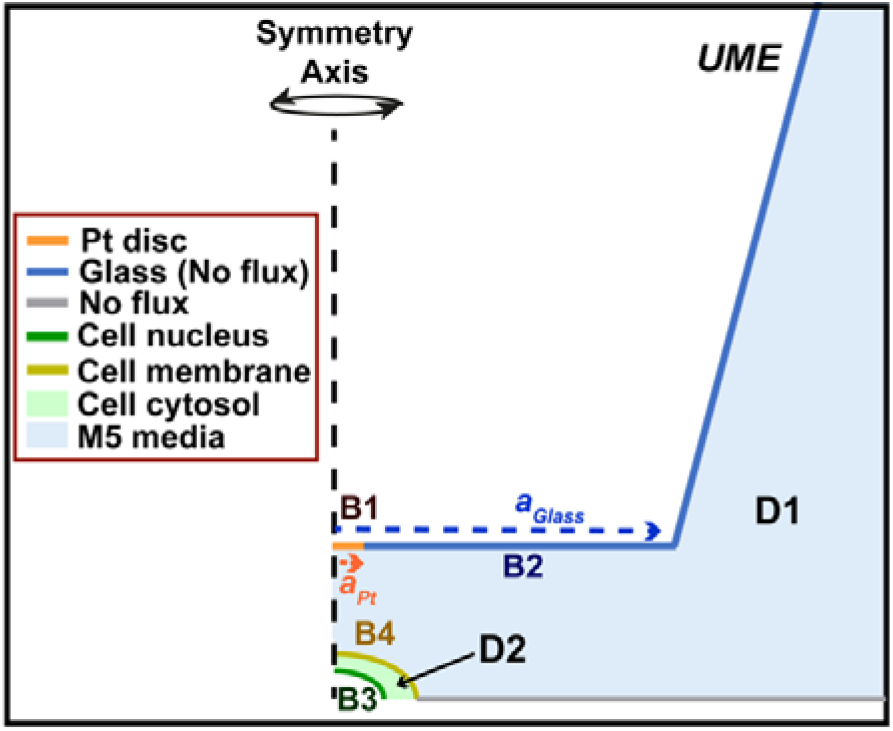
Illustrative COMSOL model, describing the domains and boundary conditions. See text and SI section SI-4, for descriptions of the different boundaries labelled B1-B4 and domains D1 and D2.

Details of the FEM model are given in the SI (Section SI-4). In brief, the “Transport of Diluted Species” COMSOL module was used to solve for oxygen, considering mass transport by diffusion only in an axisymmetric cylindrical geometry (UME directly over the center of the cell). All simulations were performed as time-dependent studies, for pulse lengths equivalent to those performed in the SECM experiments (1 second). With reference to Figure 2 and Table S1 (SI), the boundary condition, B1, represents the electrode, where oxygen is reduced at a sufficiently fast rate to ensure diffusion-limited conditions, and resulting in the concentration of oxygen at the UME surface being (close to) 0. B2 represents the glass sheath, and was therefore set as a no-flux boundary. Regarding the cell, B3 represents the outline of the cell nucleus, where the rate of cell oxygen consumption is set by a 1^st^ order rate law (see section SI-4), and varied over a set of values. B4 represents the cell outer membrane, which demarcates a change in the diffusion properties of O_2_ (but with no kinetic barrier to transport between the cell and bathing solution). The UME presence in close vicinity of the cell will decrease the oxygen concentration at the cell domain (D2). We opted to keep OCR values constant by increasing the kinetic constant. Our calculations therefore represent a best-case scenario for examining SECM measurements of OCR, by maintaining the value irrespective of the action of the UME.

## Results and Discussion

### Self-referencing SECM measurements of single cell OCR

As a proof-of-concept of the self-referencing SECM method we measured single cell OCR of HeLa cells. We combined this approach with CLSM, which allowed the use of tetramethylrhodamine, methyl ester (TMRM), an indicator of mitochondrial membrane potential,^48–50^ which is a key bioenergetic variable relating to cell respiration rate.

Figure 3 demonstrates the power of self-referencing SECM for an experiment that consisted of taking 1 second ORR pulse measurements in the vicinity of a cell (10 μm from the cell surface) and in the bulk, repeated 30 times (with 30 seconds interval between each measurement cycle). Figure 3A shows the near cell (surface) and bulk ORR current over time, each data point is an average of the last 20 data points, (over ca. 20 ms duration) of the 1 second ORR pulse measurement. The absolute value of the near-surface currents decreases over time, and a similar trend is also observed for measurements in the bulk (Figure 3A, blue trace), where the oxygen concentration is stable. This deterioration of response is attributed to electrode fouling that evidently has a very significant effect on the UME current response over time; such effects are rarely considered in single-cell SECM measurements.

**Figure 3.**
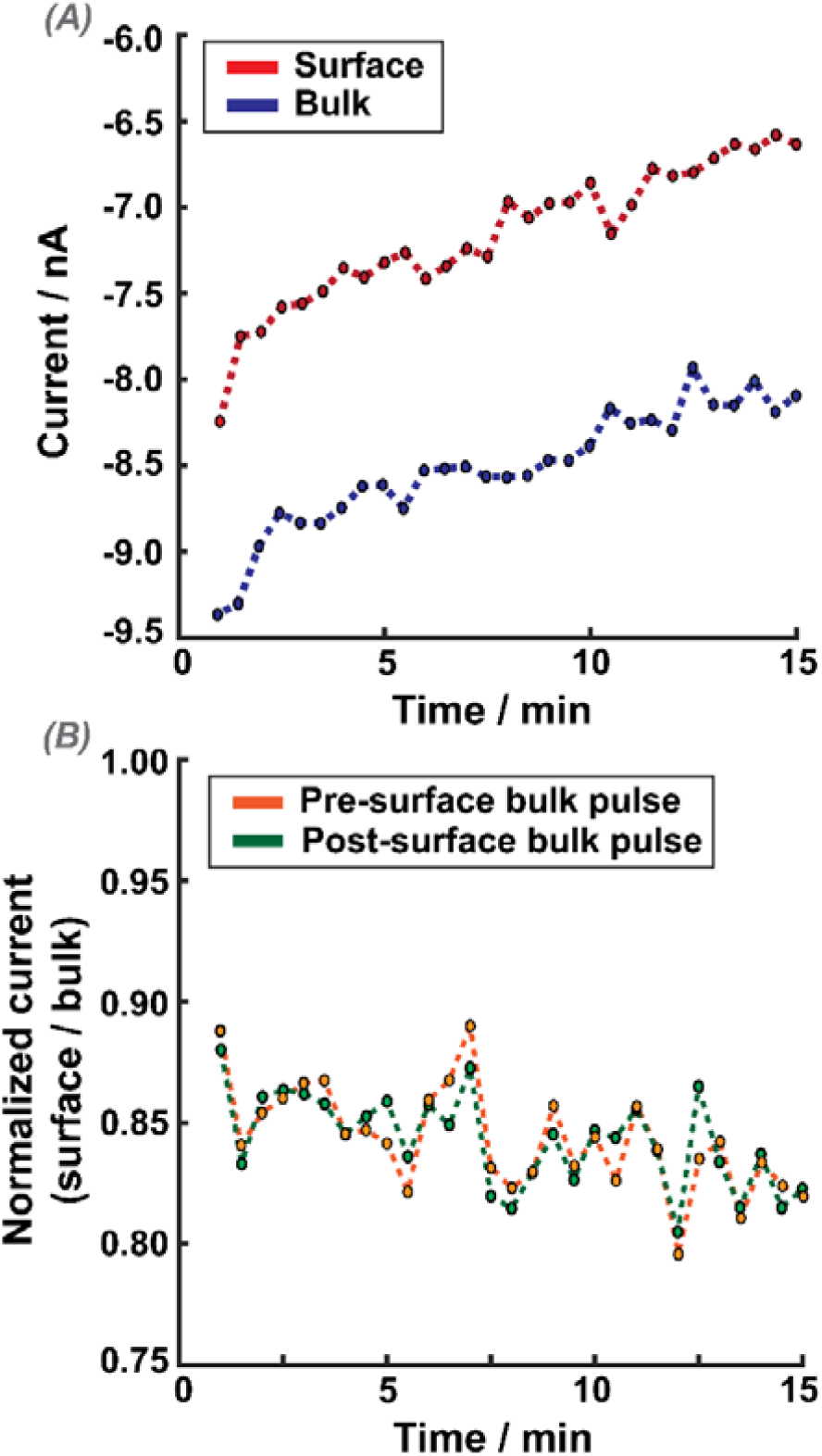
Experimental results from a self-referencing SECM experiment. (A) The current recorded for each OCR measurement in the bulk (blue) and at the cell vicinity (near-surface) (red). (B) Near-surface current normalized to the current measured in the bulk, either before the surface pulse (orange), or immediately after the surface pulse (green). 30 repeated measurement cycles, each of 1 second duration, and one approach (hop) every 30 seconds.

Figure 3B shows the near-surface current normalized with respect to either the bulk current measurement preceding the surface pulse (orange trace), or the bulk measurement directly after the surface pulse (green), as a function of time. The traces are similar and normalized currents fluctuate around 0.84 (± 0.03), which is the normalized current value expected for hindered diffusion for a UME of this geometry for the defined measurement time and approach distance (*d*) *ca*. 10 μm (see Section SI-2). Indeed, a very similar trend in the current is also seen when these measurements are made directly over glass at a distance of *ca*. 10 μm, as shown in SI, Figure S-1. These experiments thus demonstrate that the measurements are rather insensitive to single-cell respiration, and the current response is predominantly due to mass transport hindrance into the UME/surface gap.

### Does increasing the OCR allow single-cell detection by SECM?

To determine if it was possible to detect a definitive signal for cell respiration using SECM, measurements were taken before and after the addition of carbonyl cyanide chlorophenylhydrazone (CCCP, 2 μM). CCCP is an ionophore that uncouples respiration from oxidative phosphorylation, by collapsing the proton motive force at the mitochondrial membrane, and is shown to reduce membrane potential and increase cellular OCR values.^1, 51, 52^ To verify the impact of CCCP, the cells were additionally stained with TMRM, a commonly used membrane potential indicator, and visualized by CLSM during the SECM measurements. As above, SECM current was sampled for 1 second both near the cell and in bulk (self-referencing protocol described). This procedure was repeated 50 times with an interval of 30 seconds between each of the near-cell measurements. Figure 4 shows a typical response in terms of raw current in the bulk and near-surface (Figure 4A), the normalized current (Figure 4B), and the TMRM mean intensity at the cell and background (Figure 4C and 4D); 2 regions for each. CCCP was added to the bulk solution at the 10-minute mark.

**Figure 4.**
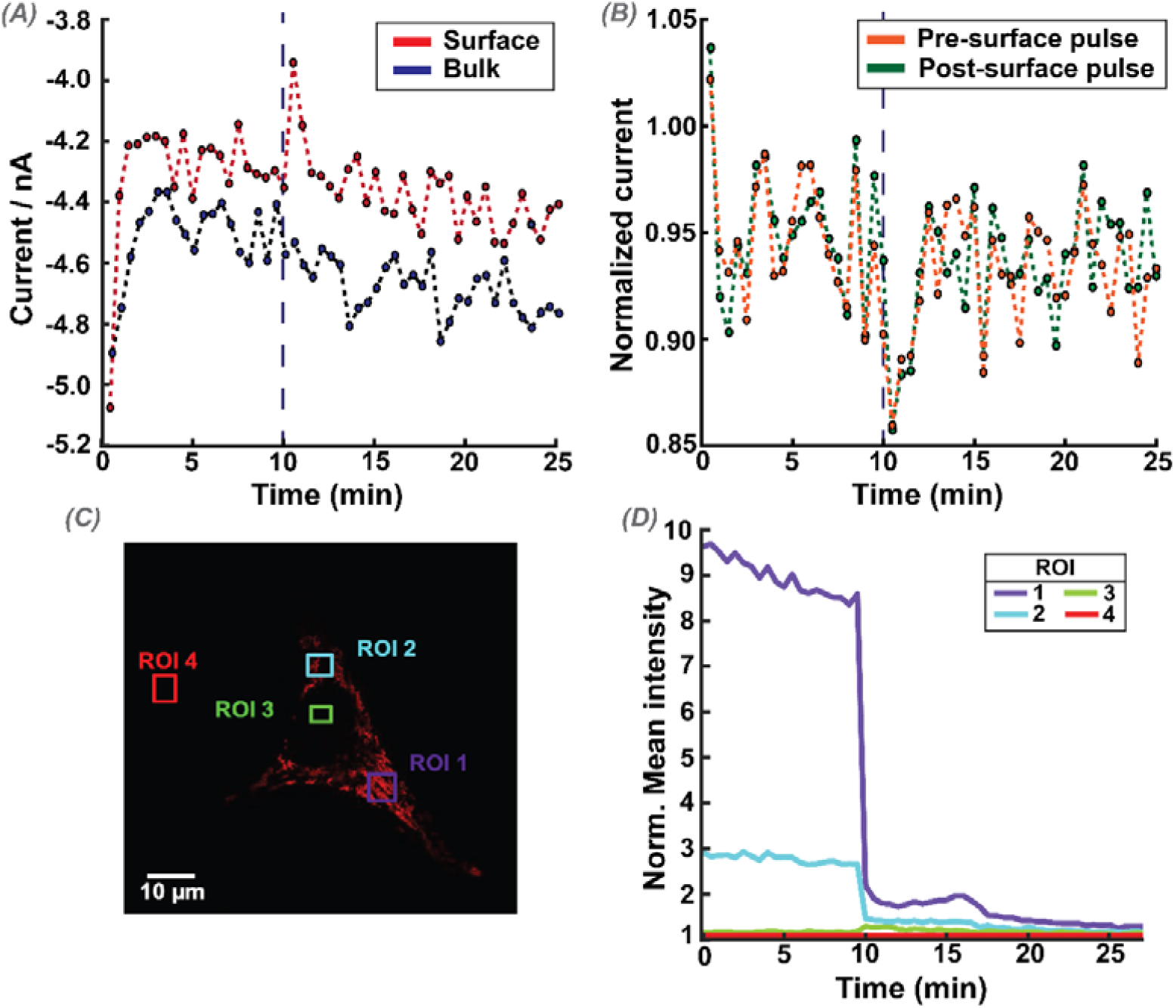
Self-referencing SECM measurements of OCR at an HeLa cell, with the addition of CCCP (2 μM) at 10 minutes-indicated with a grey dashed line. (A) The raw bulk (blue) and surface (red) currents, with each point as an average of the last 20 points at the end of each pulse. (B) Near-cell current normalized to the current measured with the bulk pulse, either before the surface pulse (orange), or after the surface pulse (green). (C) CLSM image of the cell under study, stained with 50 nM TMRM. (D) Each colored line shows the mean fluorescence intensity for a given region of interest (ROI, drawn boxes with the same colors), normalized to the background (ROI 4).

Before addition of CCCP, using either the pre-surface or post-surface bulk pulse as the reference signal resulted in no significant difference in the normalized currents compared to expectations for an inert surface. As can be observed, the TMRM fluorescence across the cell was immediately reduced to near background levels upon the addition CCCP, indicating the mitochondria membrane potential collapse as shown before.^50^ Within the variability of these time-course measurements, there is no detectable difference in the normalized current response before and after TMRM. This contrasts with the stable increase in OCR seen in population-level measurements upon application of CCCP.^1, 51^ As TMRM is a reversible reporter,^53, 54^ any possible recovery of oxidative phosphorylation coupling to respiration can be disregarded as there is no return in TMRM fluorescence. We can reasonably assume that the cell would have a higher OCR during the entire period of this experiment after the addition of CCCP, but this goes undetected by self-referencing SECM.

### FEM modeling highlights inherent limitations of SECM for the measurement of single-cell OCR

To better understand any limitation of an SECM-based OCR measurement, we developed an FEM model to mimic the experimental setup used here (see *Methods* and SI-4). In brief, we simulate ORR at an UME that is immersed in an aerated bulk medium and set an OCR at a targeted cell. Figure 5 presents simulation results from this model, for a 5 μm radius UME (a commonly used size in SECM experiments and in this work),^1, 55^ where ORR is diffusion-controlled. We simulate two scenarios, where the UME is held at 100 μm (Figure 5A) and 10 μm (Figure 5B) over a single cell consuming oxygen at 1×10^−11^ mol s^-1^. This rate is used for illustrative purposes and is *ca*. 10^6^ times faster than previously reported population-based single cell OCR values for HeLa of 1×10^−17^ mol s^-1^.^31, 56^

**Figure 5.**
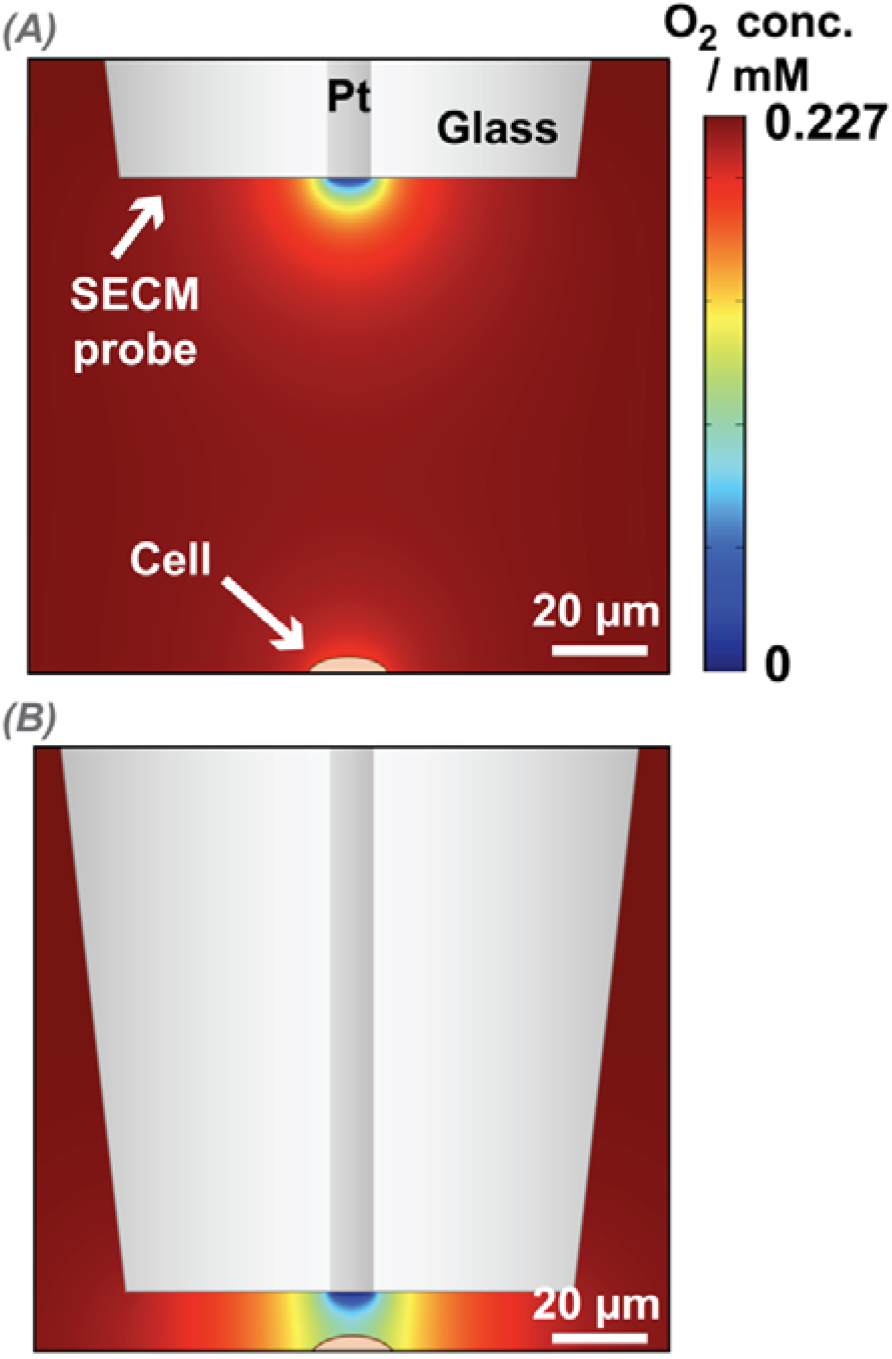
FEM-based simulation of single cell OCR and SECM-based ORR, with the SECM probe (platinum UME, a = 5 μm) reducing oxygen at transport-limited rate, and an illustrative cell – orange semi ellipsoid-undergoing oxygen consumption (for illustration, OCR = 1×10^−11^ mol s^-1^). The color gradient depicts the oxygen concentration in solution, with the deepest red at the top of the scale representing the bulk oxygen concentration (0.227 mM).^59^ (A) SECM probe positioned 100 μm above the cell (effectively in bulk position). (B) SECM probe held 10 μm above the cell.

The simulation results shown in Figure 5 highlight that ORR at a platinum UME results in a much stronger oxygen sink than OCR at the cell undergoing respiration. This is significant because for detectable measurements the cell needs to ‘shield’ the UME from part of the oxygen diffusional flux, as discussed before in the context of SECM ‘shielding’ (or, equivalently, ‘redox competition’) measurements with oxygen detection.^57, 58^

To measure the OCR of a cell, the SECM probe must be affected by OCR at the cell which competes with the tip for oxygen.^58^ From the FEM simulations we can calculate how far the oxygen gradient extends from the cell surface into bulk solution for different OCR (without the presence of the UME). Based on the OCR of a single HeLa cell being about 1×10^−17^ mol s^-1^,^31, 56^ OCR values were explored over several orders of magnitude around this value. The concentration profiles extending from the cell surface into solution, normalized by bulk oxygen concentration, are depicted in Figure 6. At an OCR of 1×10^−16^ mol s^-1^, the oxygen gradient is very shallow, with oxygen levels only changing by 0.23 % within 5 μm from the cell at the axis of symmetry. The situation is similar for other OCR values considered, with rates at and less than 1×10^−16^ mol s^-1^ showing little change of oxygen concentration even on a sub-micron scale from the surface (see Figure 6 inset).

**Figure 6.**
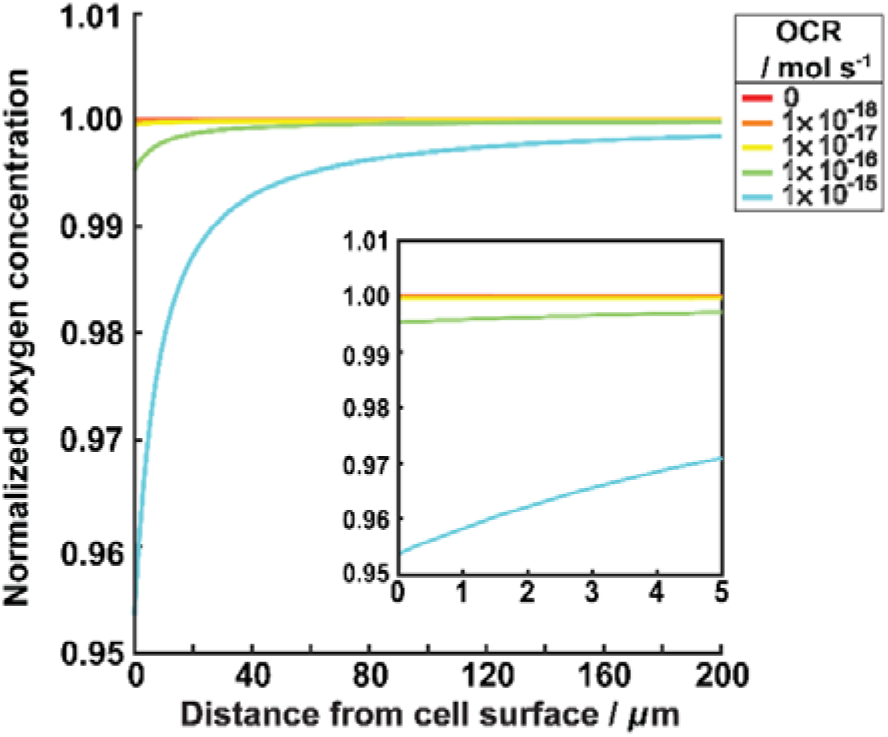
FEM simulation of the local oxygen concentration normalized to the bulk concentration, as the distance between measurement point and the cell surface is increased (x-axis, with 0 *μ*m indicating measurement at cell surface). Differently colored lines show results for different OCR values (mol s^-1^) as shown on the key. The inset shows a magnified section of the x-axis plot. Green line is the closest to the reported single-cell OCR of HeLa cells (1×10^−17^ mol s^-1^).^31^

### Effect of UME-cell distance directly impacts on SECM-based OCR measurements

We explored whether a smaller UME would be more sensitive to OCR, noting that it would need to be positioned at closer distances to the cell when operated in the shielding mode of detection. We simulated the normalized current response of a 1 μm radius UME (RG = 10) for different UME-cell distances and for different cellular OCR values. The UME-cell distances (*d*), are normalized to the UME radius (*a*), *i*.*e. d/a*. In Figure 7, all currents are normalized to the case of OCR = 0 mol s^-1^ (no oxygen consumption), thereby accounting for any effects arising from hindered diffusion due to close UME-cell proximity.^60^

**Figure 7.**
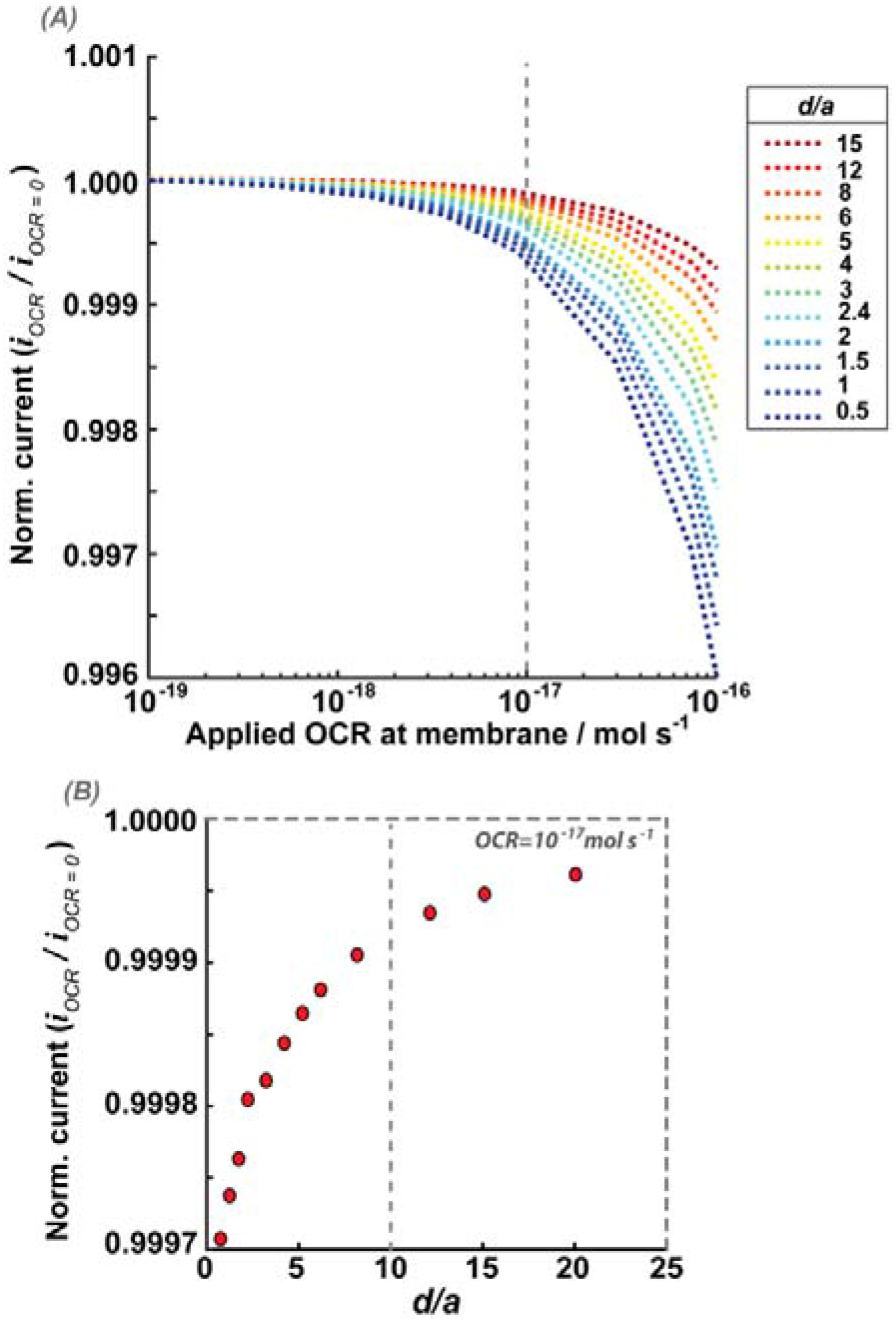
Effect of UME-cell distance on current measurements. The normalized currents are simulated for a UME performing ORR at a transport-limited rate and a cell respiring at different OCR values. Currents are normalized by the UME current at the same height (UME-cell distance) with OCR set to 0 mol s^-1^. (A) Normalized current against cellular OCR. Each colored line shows current for a given UME-cell normalized working distance (d/a). Distances are normalized to the UME radius of 1 *μ*m. Dashed line marks the simulated OCR condition that is closest to the reported OCR of HeLa cells (1×10^−17^ mol s^-1^).^31, 56^ (B) Normalized current against UME-cell normalized working distance at the simulated OCR condition that is closest to the reported OCR of HeLa cells (1×10^−17^ mol cell^-1^ s^-1^) – the dashed line in panel (A). The dashed line in panel (B) marks the UME-cell normalized working distance of 10 (UME stationed 10 *μ*m from the cell surface).

From Figure 7 we find that for smaller UME-cell distances, there is a larger decrease in current magnitude for a given OCR. This is expected, due to the larger intersection between the two diffusion layers (UME and cell), resulting in the UME ‘sensing’ more of the cellular OCR. For an OCR of 1×10^−17^ mol s^-1^, which is within the range of reported OCR for most cells,^25, 32^ the SECM probe would need to be at a working distance of closer than 8 μm from the cell to record a normalized current difference between a respiring and non-respiring cell of > 0.0001 (0.066 pA). This would require extraordinary measurement precision. For instance, for a UME with 1 μm radius and UME-cell distance of 1 μm, the change between a non-respiring and a cell respiring at a rate of 1×10^−17^ mol s^-1^ is only 0.157 pA, with respect to a baseline of 663.8 pA for the non-respiring cell. Measuring such small differences in current values, even if ultra-precise positioning of the SECM probe could be achieved, borders on the impossible in any environment, let alone cell media.^61^

It is clear that smaller UME-cell distances will increase the OCR measurement sensitivity, but there is a limit as to how close the UME should be to the cell, without undue influence from the effect of SECM-induced O_2_ transfer from the cell by the action of the UME.^62, 63^ For an UME of 1 μm radius and at a distance of 1 μm from the cell, as deduced above, the oxygen flux towards the UME is *ca*. 1.72×10^−15^ mol s^-1^ (based on a current of 664 pA, resulting from simulations shown in Figure 7). This is 2 orders of magnitude larger than the cell OCR value of 1×10^−17^ mol s^-1^, demonstrating that in this experimental setup, the SECM-based measurement significantly depletes the surrounding cell environment of oxygen, and induces oxygen transfer from the cell to the UME, effectively making the cell an oxygen source to the UME (*vide infra*).^29, 62, 64^

To further explore this aspect of the effect of the UME and the balance between detecting OCR and inducing oxygen transfer out from the intracellular region, we simulated the oxygen flux over the cell membrane of a single cell, during SECM-based OCR measurements using a hypothetical 1 μm radius UME (Figure 8).

**Figure 8.**
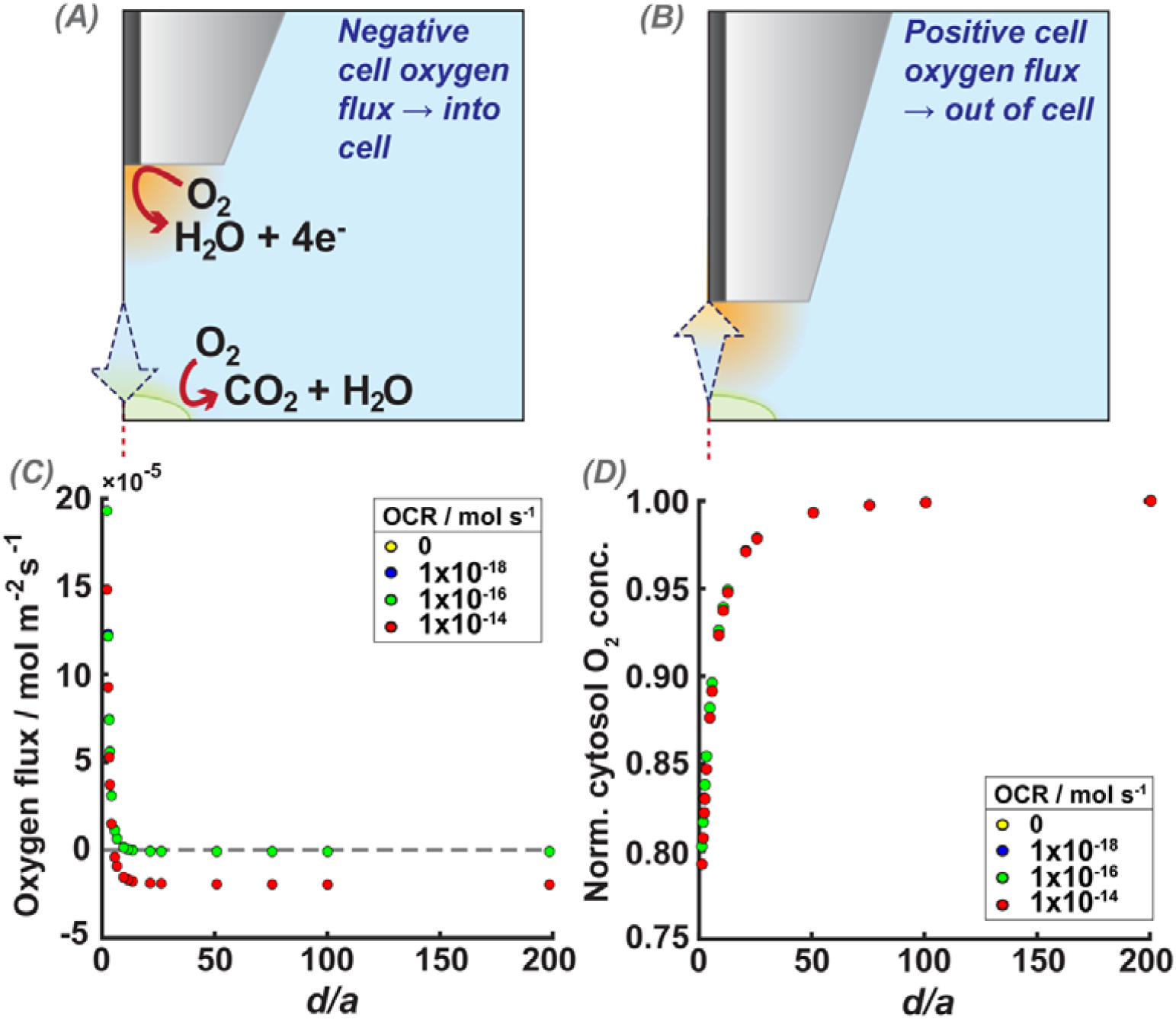
Effect of the UME on the oxygen flux at the cell/solution interface. Flux in the z-axis is used to represent the net oxygen flux direction with respect to the cell surface. (A & B) Cartoon representation of expected direction of oxygen flux when the UME is far away in bulk (negative flux values), and when the UME is close enough to the cell, to impact the cell oxygen flux (positive flux values). (C) Simulated z-direction oxygen flux at the top of the cell for different normalized UME-cell distances (d/a), for different OCR values. (D) Simulated oxygen concentrations in the cell cytosol (see Figure 2, D2) normalized to the intracellular oxygen concentration when the UME is sufficiently in bulk and inactive to impact the cell, for different normalized UME-cell distances (x-axis, d/a), where the cell is respiring at different OCR rates. In (C) and (D), the OCR of 0 and 1×10^−18^ mol s^-1^ (yellow and blue) are completely overlapped by the 1×10^−16^ mol s^-1^ series (green).

The oxygen flux perpendicular to the cell membrane (*z*-axis) was measured at the top of the cell. During respiration, oxygen is transported from the media into the cell, resulting in a negative flux (as defined in the model herein) across the membrane. When the UME is brought into close proximity to the cell, the UME ORR reaction may induce oxygen transport out of the cell towards the electrode (Figure 8B). In Figure 8C and 8D, the data for the OCR being 0 and 1×10^−18^ mol s^-1^ (yellow and blue) cases overlap with those for the 1×10^−16^ mol s^-1^ case, demonstrating how close this range of OCR is to a non-respiring cell. For UME-cell separations less than 12 μm the overall flux direction is out of the cell (induced transfer) and the intracellular oxygen concentration decreased monotonically at increasingly smaller distances (see Figure 8D). This shows clearly that SECM measurements can readily induce hypoxic cell conditions.^62, 63^

The simulations imply that there would be advantages to decreasing the UME flux, so as not to overwhelm the cell respiration process. This could be achieved by reducing the applied potential to the UME. However, operating the tip under kinetic control would be very difficult in a biological medium, where electrode fouling is problematic. Another alternative method could be to coat the UME surface in a polymer layer with the aim of slowing down the diffusion of oxygen at the UME,^65, 66^ although care would be needed to ensure oxygen did not have high solubility in the membrane, so that it would act as an oxygen sink.

### Is there an ideal UME size and SECM working distance for single cell OCR?

We now consider if even smaller electrodes would be useful for single-cell OCR measurements, noting that nanoscale UMEs have been deployed previously for live cell SECM measurements.^1, 21, 67^ Figure 9A shows the effect of normalized working distance on UME normalized current (with respect to an inert surface at the same UME-surface distance) over a cell with an OCR of 1×10^−16^ mol s^-1^ (an order of magnitude larger than the typical value; *vide supra*) for an UME radius of 0.25, 0.5 and 1 μm. The biggest difference in normalized current between a cell undergoing OCR and the baseline inert surface response is found with the smallest UME radius (0.25 μm), although the effect of OCR on the UME current is still close to negligible even at a very small normalized distance.

**Figure 9.**
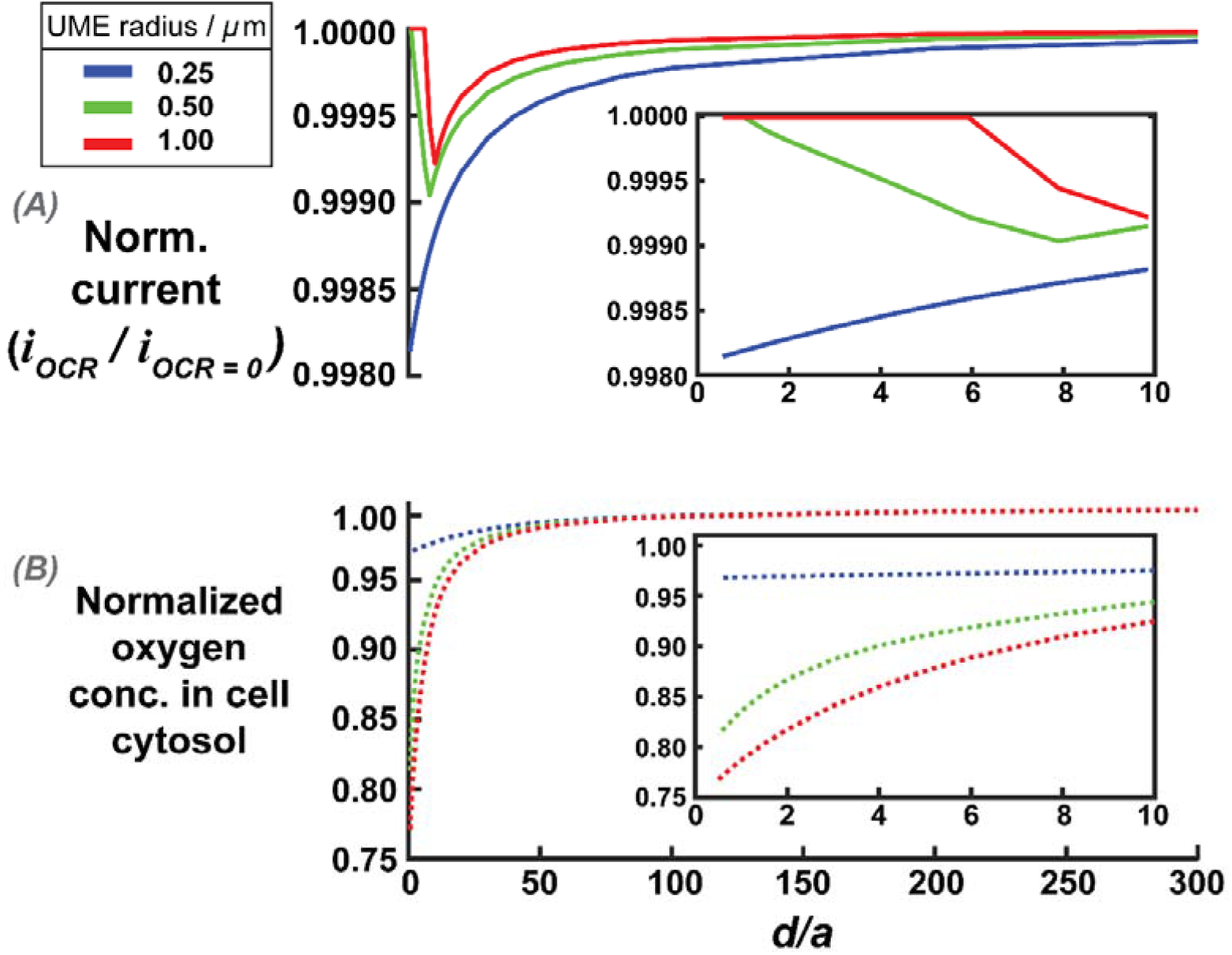
(A) Simulated normalized currents against normalized UME-cell distance (d/a) for different UME radii (as defined in the key). The current is normalized to that for a non-respiring cell at the same distance. The inset highlights the normalized currents for normalized distances between 0 and 10. (B) The average oxygen concentration in the cell cytosol as a function of normalized UME-cell distance. Concentrations are normalized to the cytosol concentration when the UME but is not active. The inset highlights the normalized oxygen concentration for distances between 0 and 10.

The effect of SECM induced oxygen transfer from the cell again decreases the cell cytosol oxygen concentration. Figure 9B shows the oxygen concentration inside the cell, normalized to the concentration where the UME is not active (no electrode reaction), for the different UME radii. For the smallest electrode radius, 0.25 μm, the UME induces oxygen transfer from inside the cell at normalized working distances smaller than 0.5, when the cytosol oxygen concentration decreases by 4 %. Oxygen transfer from the cell is also evident at the normalized current profiles in Figure 9A, where there is an increase in normalized current for the 0.5 and 1 μm radius cases when the UME-cell distance becomes sufficiently small. This is again due to the cell acting as a local source of oxygen for the UME reaction (induced transfer).^63^

These simulation results demonstrate the stringent requirements towards optimal SECM conditions for measuring cellular OCR; but they highlight major practical challenges. In particular, biofouling is more problematic with the use of sub-micron, or nanoscale, UMEs makes (rates of diffusion to UME proportional to inverse of UME radius).^68^ Furthermore for a normalized distance of 0.5, a 0.25 μm radius UME would need to be placed 0.125 μm above the cell with high precision. Even accepting an oxygen loss of 4 % from the cell, the UME would record a normalized current of *approx*. 99.8 % of that for an inert surface, for a cell respiring at 1×10^−16^ mol s^-1^ (an order of magnitude higher than typical, *vide supra*). These measurements are thus highly challenging.

### Conclusions and Perspective

By combining experimental self-referencing SECM measurements with FEM simulations we have elucidated significant limitations of single-cell OCR measurements. Our experimental results have revealed that even when self-referencing SECM is used (which is an improvement on conventional SECM by using the updated bulk signal throughout a measurement sequence), it is highly difficult, if not possible, to detect OCR practically. Furthermore, by tracking the bulk UME response over time in these measurements, we showed that the UME response deteriorated significantly, which would greatly impact conventional SECM measurements. Our simulations, exploring a range of SECM conditions, have revealed that it is extremely challenging to measure the OCR at a single cell due to the small rates and consequent tiny oxygen gradients that result. In essence, the major challenge for single-cell OCR measurements with SECM is: how can such small oxygen fluxes be measured without disturbing the cell function?

The optimal conditions for self-referencing SECM measurement of cellular OCR result from the use of sub-micron sized SECM probes, which allow closer working distances to the cell, with increased sensitivity, but without much disturbance to the cell physiology. These experimental conditions are recognizably challenging, in particular with regard to the high precision that would be required in the approach distance and in nanoelectrode fabrication. To date, most studies involving SECM and cells have used UMEs ranging from 0.5 μm to 10 μm in radius. Although such large UMEs can be used for some SECM-based flux measurements, without self-referencing and the aid of FEM simulations, previous reports of OCR measurements should be considered semi-quantitative. As a result of the work herein, we suggest an experimental framework for single-cell SECM OCR measurements to reduce perturbation of the cell conditions by SECM and to account for electrode fouling for long duration measurements.

We note that our experimental work focusses on HeLa cells, which are known to have a reduced respiration rate due to their preference for glycolytic metabolism,^69^ therefore practically making it more difficult to experimentally probe for respiration. However, our FEM models were extended to cover a wide range of the respiration rates, covering OCR rates founds across a range of different commonly studied cell lines, and extending beyond the expected range for single cells (1×10^−16^ mol s^-1^ used herein for many calculations is higher than reported for any single cell based on bulk measurements). Future studies can explore the same measurement for different, more respiratory cell lines or for groups of cells.

The combination of SECM, fluorescence microscopy and FEM simulations offers exciting possibilities for future work, where the SECM probe can be used as an actuator to shift cell homeostasis in a controlled manner (supported by FEM) and fluorescent dyes are used as quantitative reporters of the cellular status. Furthermore, FEM models can be increasingly tailored to better represent the biochemical and physical properties of a cell,^70^ with the possibility of full 3D architecture to best account for unique cell geometry.

## Author Contributions

Methods were designed by KC, DV and GNM, with experiments performed by KC. Simulations were developed by KC, GNM and PRU, and performed by KC and GNM. PRU and OSS conceived the study, supervised and helped with data interpretations. The manuscript was written by all authors.

## Acknowledgements

KC would like to thank the EPSRC for support through MAS CDT, grant number EP/L015307/1. GNM acknowledges support from the European Union’s Horizon 2020 research and innovation program under the Marie Skłodowska-Curie grant agreement 790615 (FUNNANO). PRU thanks the Royal Society for support through a Wolfson Research Merit Award. The authors would all like to acknowledge the support of the Bio-Electrical Engineering Innovation Hub, University of Warwick, funded by the UK’s Biological and Biotechnological Sciences (grant no. BB/S506783/1) and Engineering and Physical Sciences Research Councils.

