## supplementary information for "Can Single Cell Respiration be Measured by Scanning Electrochemical Microscopy (SECM)?"

### SI-1 HeLa cell culture and media preparation

HeLa cells were obtained from Public Health England, supplied by the European Collection of Authenticated Cell Cultures (ECACC, catalogue number: 93021013). Cells were grown in Minimum Essential Medium Eagle (Sigma Aldrich, M2279) supplemented with L-glutamine (used in 100-fold dilution, Sigma Aldrich, G7513), penicillin and streptomycin (used in 1000-fold dilution, Sigma Aldrich, P4333), heat-inactivated fetal calf serum (10-fold dilution, HIFC, Sigma Aldrich, 12106C), and non-essential amino acids (100-fold dilution, Sigma Aldrich, M7145). Due to the sodium bicarbonate buffer used, cells were incubated at 37 °C in 5% CO_2_. Cell flasks were passaged when the confluency reached approximately 80% (no more than a week), and never exceeded a passage number of 10.

Individual sample dishes of cells were prepared on 50 mm WillCo Wells (Glass thickness No. 1.5, USE, HBST-5040) the evening prior to experiments, cultures were diluted to approximately 5×10^6^ cell mL^-1^.

For the purposes of all SECM measurements, the medium was replaced with a Minimum Essential Medium (Sigma Aldrich, 56416C, referred to here as M5) buffered with HEPES (100-fold dilution, Sigma Aldrich, H0887), warmed to 37 °C.

### SI-2 SECM instrumentation

*Fabrication of SECM probes*

Disk shaped platinum UMEs with a radius, *a*, of 5 μm were prepared as described elsewhere.^1^ In short, this method involved encapsulating platinum wire (Goodfellow, 99.99 %, 10 μm diameter) in borosilicate glass capillaries (GC200-10, Clark Electromedical Instruments), which was then heated under vacuum and pulled to produce a tapered end using a PB-7 Narishige micropipette puller. The tapered end was then polished to create a smooth flat surface, and the glass sheath is receded through polishing the UME at an angle to create the desired ratio of active electrode surface (Pt) to glass (RG value) of 15. The UME surface was platinised in a 0.1 M hexachloroplatinate solution (0.1 V/s deposition). This was done, as several previous studies have reported increasing sensitivity through platinization for detection (reduction) of oxygen and peroxy species.^2, 3^ The amount of platinum deposited in the platinization was limited in order to ensure that the electrode geometry remained as a disk.

*SECM control measurements*

Figure S-1 shows the raw and normalized current response when the probe was approached to a glass slide (substrate) with the same feedback threshold as used for the cellular measurements. The normalized current fluctuates around a value of 0.85 as found for the cell measurements.

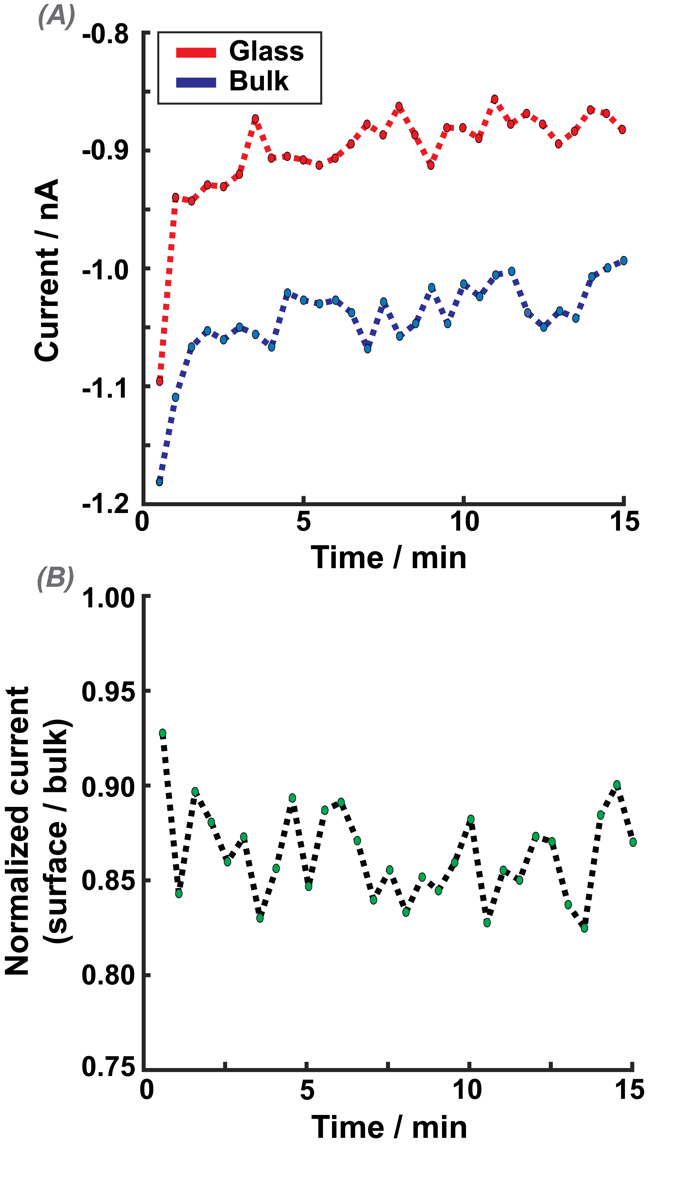

***Figure S-1*** *Experimental results from a self-referenced SECM experiment to a glass substrate. (A) The current recorded for each OCR measurement for each hop at the bulk (blue), and to a glass surface (red). (B) Surface current normalized to the current measured at the bulk position. 30 hops of each 1 second duration, 1 hop every 30 seconds.*

### SI-3 Cell staining and imaging

*Cell Staining*

Some cells were stained with Tetramethylrhodamine, Methyl Ester (TMRM, Thermo Scientific, M20036), to observe the mitochondrial membrane potential using the confocal microscope. For staining, the media was removed and replaced with PBS, and supplemented with 50 nM TMRM (prepared from a stock solution of 7.5 μM in DMSO). Cells were then incubated with the dye for 30 minutes at 37 °C, before being washed with 3×1 mL applications of sterile PBS to remove of residual dye. Fresh M5 media was then added to the plate for scanning.

*Confocal Laser Scanning Microscopy (CLSM) Imaging Conditions*

A Leica TCS SP5 X CLSM was used for imaging HeLa cells stained with TMRM, located in the mitochondrial membrane. Cells were imaged using an Argon laser, set to a wavelength of 514 nm. Emission was detected between 590 and 605 nm. A ×40 oil immersion objective was used, with the confocal pinhole set to 67.97 μm giving a section thickness of approximately 0.967 μm. The 514 nm laser power under these conditions was recorded to be 0.05 mW at the sample plane. Cells were typically imaged at 30 second intervals for the duration of the SECM experiment.

### SI-4 Finite Element Method (FEM) Simulations

*FEM Simulations*

All FEM simulations were performed with COMSOL Multiphysics (v 5.5.) using the Transport of Diluted Species module. A 2D axisymmetric simulation domain was used, as depicted in Figure 2 of the main manuscript. Mesh density (> than 35000 elements) and domain size (radial length and height of the domain > than 100*a*) were set so the simulation results are independent of both. All boundary/domain conditions are summarised in Table S-1, and simulated parameters values in Table S-2.

***Table S-1.*** *Boundary/domain conditions implemented in the FEM simulations, as shown in Figure 2 of the main manuscript. Only fluxes normal to the boundary were considered, noted by* $“\boldsymbol{n}”$*.*

| **Labelled environment (as seen in Figure 2)** | **Concentration/flux condition** |
| --- | --- |
| B1 | $\boldsymbol{n}.J_{O_{2}}=-k_{O2\_UME} C_{O_{2}}$ |
| B2 | $\boldsymbol{n}.J_{O_{2}}=0$ |
| B3 | $\boldsymbol{n}.J_{O_{2}}=-k_{O2\_cell} C_{O_{2}}$ |
| B4 | *N/A* |
| D1 | $C_{O_{2}}=227 \mu M$ |
| D2 | $C_{O_{2}}=14.5 \mu M$ |

***Table S-2.*** *Parameters used for the simulations, relating to the geometry and in other parts of the simulations.*

| **Geometry parameters** | | |
| --- | --- | --- |
| ***Symbol*** | ***Value*** | ***Description*** |
| *d* | Varied | UME-cell separation. Varied as described in the main text |
| *Cell_width_* | 9 μm | Cell width, width of B4 from origin |
| *Cell_Height_* | 2.5 μm | Cell height, height of B4 from origin |
| *Nucleus_width_* | 4 μm | ‘Nucleus’ width, width of B3 from origin |
| *Nucleus_height_* | 1.2 μm | ‘Nucleus’ height, height of B3 from origin |
| *a* | Varied | Radius of the active electrode surface. Varied as described in the main text |
| *RG* | 2 or 15 | Ratio of *a* to radius of UME glass sheath, set as 2 for *a* < 1 μm and 15 for *a >*1 μm |
| **Other parameters** | | |
| *D_O2_bulk_* | 2.2×10^-9^ m^2^ s^-1^ | Diffusion coefficient of oxygen in bulk (D1) |
| *D_O2___cyto_* | 7×10^−11^ m^2^ s^-1^ | Diffusion coefficient of oxygen in cell cytosol (D2) |
| *C_O2_bulk_* | 227 μM | Concentration of oxygen in the bulk (D1) |
| *C_O2___cyto_* | 14.5 μM | Concentration of oxygen in the cell cytosol (D2) |
| *k_O2_UME_* | 10 m s^-1^ | Rate constant for oxygen reduction at the UME |
| *k_O2_cell_* | 1.51×10^−9^ to 1.54×10^−4^ m s^-1^ | Rate constant for oxygen consumption at the cell nucleus |

Simulations consider only diffusion of oxygen (no convection), and were performed as time-dependent studies, with the duration set the same as the experiments (typically 1 second). Oxygen transport in D1 and D2 was described by Fick’s second law of diffusion (Equation S1) applied to the axisymmetric geometry of Figure 2 (main text).

$\frac{\partial C_{O_{2}}}{\partial t}=D_{O_{2}}\nabla^{2}C_{O_{2}}$	(S1)

where $D_{O_{2}}$ is the diffusion coefficient of oxygen and $C_{O_{2}}$ is the oxygen concentration.

Oxygen flux at the UME surface (B1, Table S-1) was close to diffusion-controlled owing to the large cathodic potential applied in the experiments and the oxygen concentration at the B1 is effectively 0. Oxygen flux (mol s^-1^) at the cell nucleus, simulating cell respiration, was given by a rate equation (B3, Table S-2), with the oxygen consumption rate constant (interfacial 1^st^ order kinetics), *k_O2_cell_*, varying from 1.51 × 10^−9^ to 1.54 × 10^−4^ m s^-1^. These rate constant values resulted in OCR values, calculated by integrating the oxygen molecular flux across the length of the cell wall, boundary B4 (Figure 2, main manuscript), ranging from 1.01×10^−19^ to 9.99×10^−15^ mol s^-1^. These values cover, and exceed, the range of values reported in the literature for single HeLa cell respiration rates.^4^

Simulated ORR currents were calculated by integrating the molecular oxygen flux over the UME disc (B1) and multiplying it by Faraday’s constant, *F* (96485.3 C mol^-1^), and the number of electrons, *n*, involved in the ORR reaction at platinum, reasonably assumed to be 4.^5^
